# Mechanistic Modelling of Recessive Disease through Allelic Integration of Variant Effects

**DOI:** 10.1101/2025.08.15.670494

**Authors:** Hasan Çubuk, Marcin Plech, Vahid Aslanzadeh, Marie Zikanova, Vaclava Skopova, Stanislav Kmoch, Yuxin Shen, Joseph A. Marsh, Grzegorz Kudla

**Affiliations:** MRC Human Genetics Unit, Institute of Genetics and Cancer, University of Edinburgh, Edinburgh, UK; Department of Paediatrics and Inherited Metabolic Disorders, First Faculty of Medicine, Charles University and General University Hospital in Prague, Prague, Czech Republic; School of Informatics, University of Edinburgh, Edinburgh, UK

## Abstract

Interpreting the effects of genetic variants remains a major challenge in recessive diseases, where clinical outcomes often depend on interactions between alleles. Multiplex assays of variant effects (MAVEs) measure variant function at scale, but nonlinear relationships with biochemical activity complicate the interpretation of MAVE scores. Here, we describe a broadly applicable approach to estimate enzymatic activities for thousands of genetic variants using a pair of fitness assays conducted at different expression levels, by modelling the nonlinear relationship between activity and fitness. Activity scores from two alleles are then combined into a single pathogenicity metric that captures their joint effect. We applied this approach to adenylosuccinate lyase (ADSL), a purine biosynthesis enzyme mutated in the autosomal recessive disorder ADSL deficiency. Using a yeast-based MAVE, we quantified the functional impact of over 8,000 coding variants. Our framework distinguished pathogenic from benign alleles based on estimated activity, and the integrated pathogenicity score correlated strongly with biochemical measurements from patient-derived cells, outperforming existing computational predictors. This dual innovation—the mechanistic transformation of MAVE data and allelic integration—offers a generalizable strategy for probing enzyme function and interpreting genetic variation in recessive disorders.

## Introduction

Rare diseases place a substantial burden on the healthcare system, with more than 5,000 known conditions and an estimated combined prevalence as high as 10% of the population [1]. Knowledge of genetic risk factors is key for diagnosis and treatment of rare diseases [2]. Among these factors, recessive pathogenic variants are particularly challenging to study, because they do not always show a clinical phenotype. For a recessive variant to cause symptoms, an individual must carry two mutated copies of the gene: either identical, in a homozygous state, or different, in a compound heterozygous state. Identifying such patterns can be more difficult than the discovery of dominant mutations, particularly in rare recessive diseases where patient populations are small [3–5]. Moreover, many recessive disorders arise from deficiencies in metabolic enzymes, where reduced activity leads to the accumulation of toxic intermediates or the depletion of essential products. In these cases, even partial loss of function can have severe physiological effects [6].

Difficulties faced by clinical geneticists in the identification of rare recessive mutations propagate to databases used for clinical variant interpretation. While ClinVar is an essential and widely used resource for human disease variants, it reports many variants as having conflicting interpretations or as variants of uncertain significance (VUS) [4]. gnomAD catalogues variants observed in healthy individuals, which are often considered as putatively benign [7]. This approach is effective for dominant disease, but most people carry recessive pathogenic variants in a heterozygous state [8]. Although gnomAD identifies homozygous variants, it lacks information on compound heterozygotes, limiting its utility for identifying benign variants in recessive genes.

Variant effect predictors (VEPs) are computational tools that predict the impacts of genetic variants and are widely used to prioritize those most likely to be pathogenic or benign [9]. However, VEPs show considerable heterogeneity across genes; while they perform extremely well for some genes, for others, their predictions are not much better than expected by chance [10,11]. One way of potentially overcoming the limitations of VEPs is through deep mutational scanning (DMS) and other multiplexed assays of variant effect (MAVEs), which experimentally measure the impacts of thousands of variants in parallel [12]. These approaches may be particularly useful for genes where VEPs perform poorly. We therefore identified adenylosuccinate lyase (ADSL) as a candidate for DMS: among 963 human disease genes, ADSL ranked 949^th^ in mean area under the receiver operating characteristic (AUROC) metric across 35 different VEPs for distinguishing pathogenic from putatively benign missense variants [10]. Moreover, previous studies have shown that human ADSL can complement its yeast orthologue, ADE13, offering the possibility of characterizing human ADSL variants using a yeast-based experimental model [13,14].

ADSL plays a role in de novo purine biosynthesis by converting the metabolites SAICAR into AICAR and SAMP into AMP. ADSL mutations cause the rare recessive disease ADSL deficiency. No individuals possessing biallelic null (*i.e.* nonsense or frameshift) variants have been observed to date, as complete loss of function is presumably embryonically lethal. Most clinically reported genotypes are compound heterozygotes, with 40 out of 49 unique genotypes exhibiting compound heterozygosity. ADSL deficient patients show neurological and developmental symptoms such as developmental delay, intellectual disability, seizures, autistic features, and muscle weakness [15–17].

Biochemical assays in patient samples confirm that ADSL deficiency is associated with the loss of ADSL enzymatic activity [18]. A series of double knockout experiments in yeast suggests that the physiologically important function of ADSL may lie less in purine biosynthesis, which can proceed via an alternative pathway, and more in the clearance of toxic intermediate metabolites, with SAICAR being the most likely candidate [19]. While levels of SAICAR (in its accumulated form, SAICAr) and SAMP (in its accumulated form, S-Ado) in urine, plasma and cerebrospinal fluid do not correlate with the disease severity, higher SAICAr/S-Ado ratios in cerebrospinal fluid do correlate with more severe phenotypes [16].

Here we developed a DMS assay to measure the effects of human ADSL variants using yeast complementation. We obtained fitness scores for approximately 90% of missense and stop-gain (*i.e.* nonsense) variants and then used a mathematical model to convert these into estimated enzyme activities. These estimates showed positive correlation with recombinant ADSL activity measurements. We further developed an additive model that incorporates the contributions of both alleles to the overall phenotype, combining their estimated effects into a biallelic pathogenicity score (BiPS), which correlated with biochemical ADSL activity in patient samples and improved classification of clinically reported genotypes. This approach provides a framework for interpreting recessive variants and may support more accurate diagnosis in ADSL deficiency and other disorders associated with compound heterozygosity.

## Results

### The fitness landscape of human ADSL

We designed a yeast complementation assay to investigate the effects of missense variants in human adenylosuccinate lyase (ADSL). In our assay, ADSL variants are scored for their ability to support growth of the *Saccharomyces cerevisiae* strain yTHC-ADE13, in which the homologue of ADSL, ADE13, is conditionally depleted using a doxycycline regulated promoter. ADE13 is essential for yeast survival, and loss of its function arrests growth [19]. Treatment of cells with doxycycline resulted in a complete growth arrest within 48 hours in continuous treatment. Yeast growth could be rescued by expressing ADSL from centromeric plasmids (**Fig. 1a-b**). To allow quantitative modelling of the relationship between ADSL expression levels and growth, we utilized two synthetic promoters with varying yields of ADSL. A weak promoter, XS, resulted in partial complementation, while a strong promoter, XL, fully restored growth (**Fig. 1b**). Expression of human ADSL also caused a small increase in growth in the presence of ADE13 (i.e., in the absence of doxycycline) (**Fig. 1b**), perhaps indicating that the doxycycline-regulated promoter in our strain is less effective than the original promoter.

**Figure 1:**
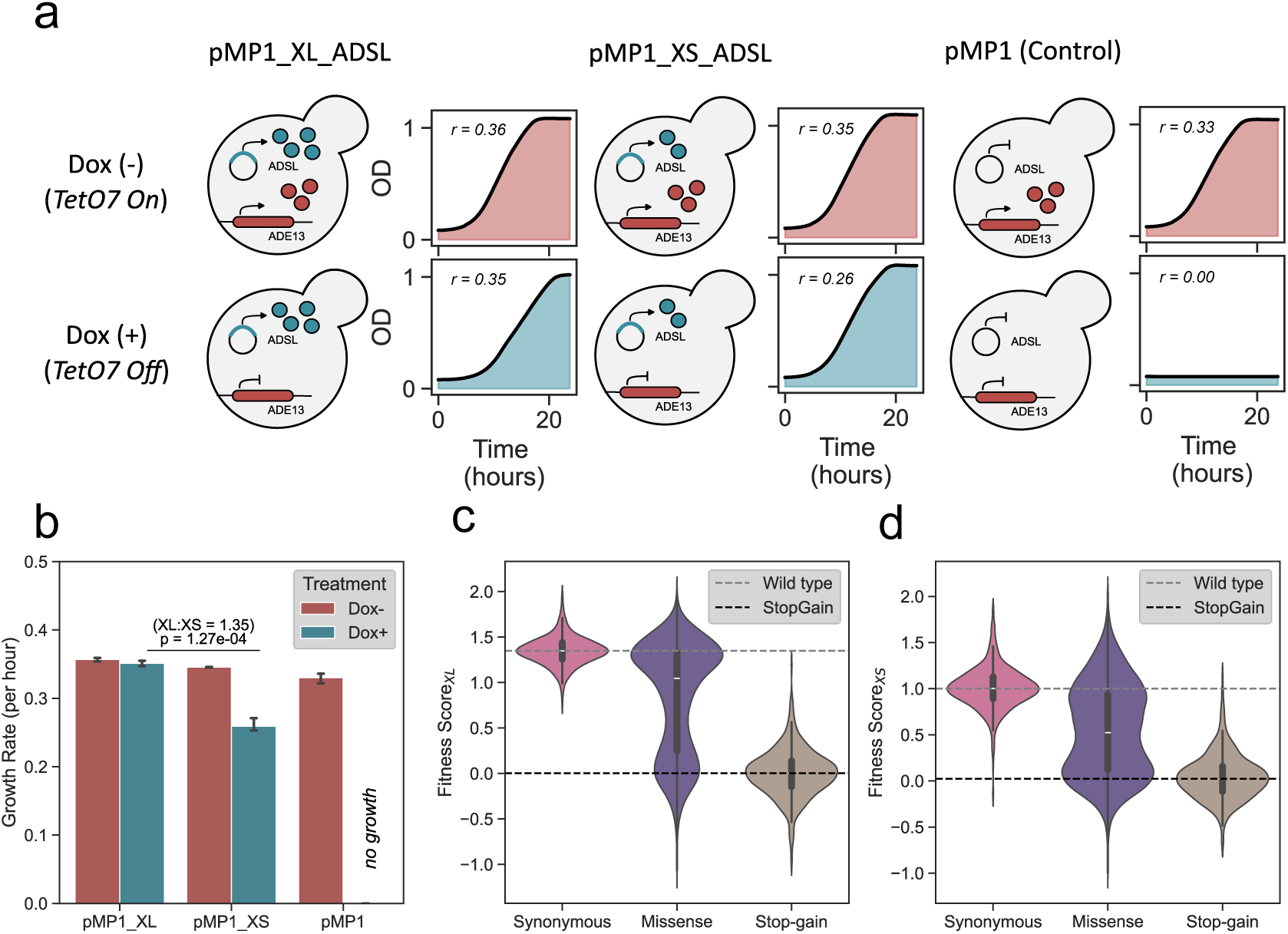
Functional complementation of yeast ADSL (ADE13) with its human homologue (ADSL) and DMS-derived fitness scores. a. Growth curves of yTHC-ADE13 strains expressing human ADSL at high (*pMP1_XL*), low (*pMP1_XS*), or no (*pMP1*) levels under doxycycline-untreated (Dox-) and doxycycline-treated (Dox+) and conditions. Dox+ suppresses ADE13 expression while untreated condition allows ADE13 expression. OD measurements were recorded over 24 hours, and each curve is the average of three replicates. r, growth rates.
b. Growth rates of the same strains in panel a under Dox+ and Dox-conditions. Each condition was measured in triplicate. Under Dox+, the growth rate for high expression was 1.35 times that of low expression (p = 1.27 × 10⁻⁴, t-test).
c. Distribution of DMS-derived fitness scores for synonymous, missense and nonsense (stop-gain) variants in high expression (*pMP1_XL*) under Dox+.
d. Same as C, but for low expression (*pMP1_XS*). Median scores for wild-type and stop-gain variants are marked with dashed lines.

We generated a library of ADSL variants in the Gateway Entry vector pAINt and transferred them to barcoded destination vectors containing either the weak (pMP1_XS) or strong (pMP1_XL) promoter. Long-read sequencing enabled phasing of barcodes with variants, resulting in coverage of 89.5% and 89.0% for missense and stop-gain variants in pMP1_XS, and 93.3% and 95.4% in pMP1_XL, respectively. Across the library, 60.5% of barcodes were associated with missense variants and 2.6% with stop-gain variants; the remainder were associated with wild-type, synonymous and multiple variants **(Supp. Fig. 1a-b)**. On average, each missense or stop-gain variant was linked to 9 barcodes in pMP1_XS and 13 in pMP1_XL.

ADSL libraries were transformed into yeast and competitive growth assays were conducted for each library for 5 days in the presence and absence of doxycycline, with samples collected every 12 hours (ten time points in total). After isolating plasmid DNA, we amplified and sequenced barcodes, and calculated barcode scores using the Enrich2 pipeline. We filtered data using internal reproducibility metrics to ensure high data quality and high coverage **(Supp. Fig. 1c-d)**. This resulted in 8605 (89.1%) missense and stop-gain variants scored in the XS experiment and 8990 (93.1%) missense and stop-gain variants in the XL experiment. To account for differences in complementation with weak and strong promoters, we normalized fitness scores so that the median score of stop-gain variants was equal to 0, and the median score of wild-type barcodes corresponded to the growth rate of the respective strain: 1.0 for XS, and 1.35 for XL experiments (see Methods). This normalization enabled direct comparison between the XS and XL experiments. We verified that normalized fitness scores showed a linear relationship with cumulative growth measured in standard optical density-based growth assays for selected ADSL missense variants **(Supp Fig. 1e**).

We next analysed the distributions of fitness scores of synonymous, missense, and stop-gain variants (**Fig. 1c-d**, **Fig. 2**). As expected, scores of stop-gain variants were tightly centred around 0, synonymous variants behaved like wild-type ADSL, and most missense variants were distributed between those of stop-gain and synonymous variants. Missense variant scores were on average lower (closer to stop-gain) in the XS experiment when compared with the XL experiment; we discuss this in detail in the next section.

**Figure 2:**
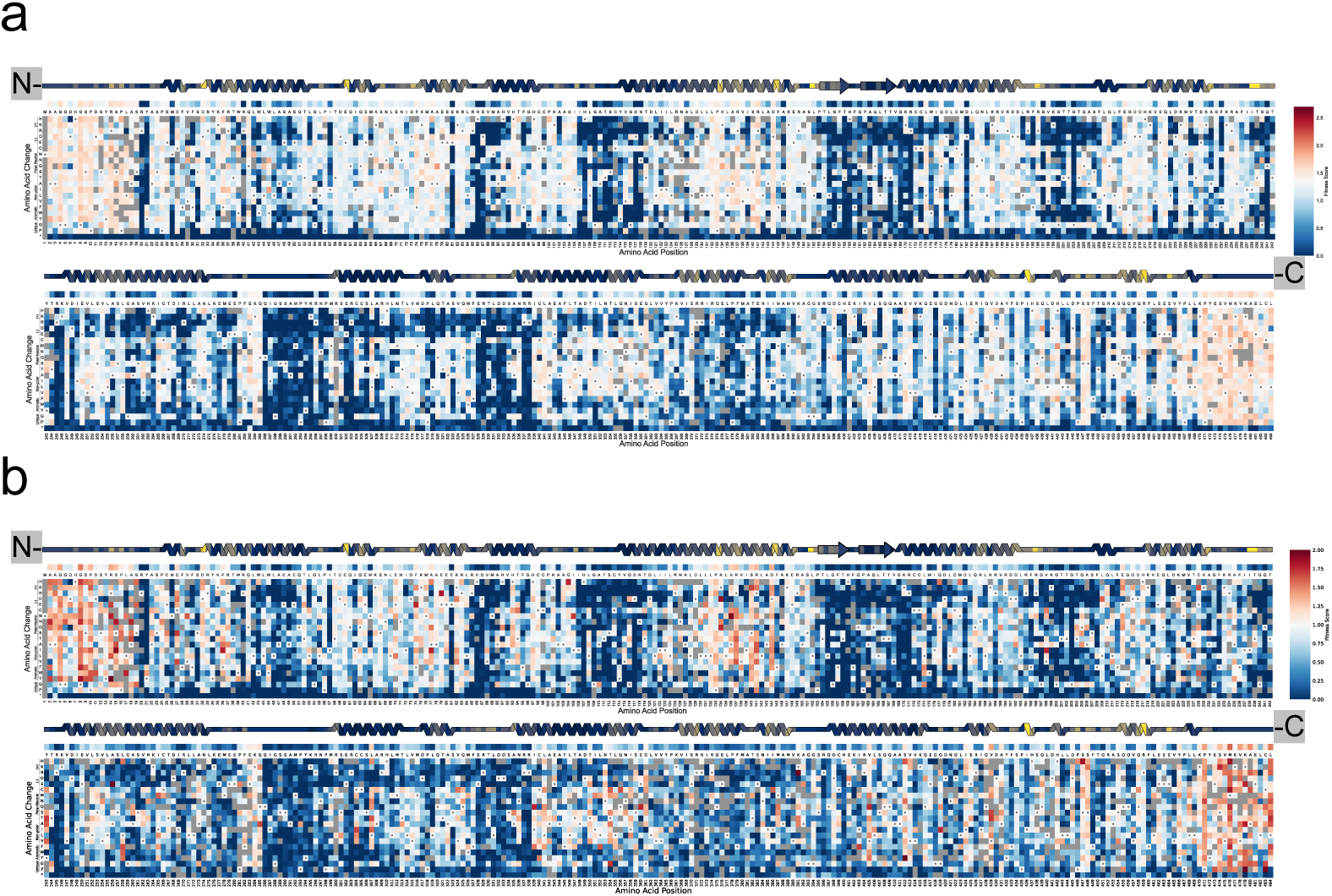
Fitness score landscape for the human ADSL gene in XL and XS experiments. a. Heatmaps of DMS-derived fitness scores in XL (a) and XS (b) under doxycycline treatment, each including: Secondary structure with residue conservation (top, viridis scale). Position-specific median fitness scores for missense variants (middle), Comprehensive fitness scores for all missense and stop-gain variants (bottom). Color scale: darker blue (damaging), white (wild-type-like), red (hyper-complementing). Damaging variants show similar patterns in both conditions. Each heatmap is split into two sections for improved visualization.

As expected, mutational effects correlated with evolutionary conservation, and mutations to prolines and to charged amino acids were typically more damaging than mutations to polar or non-polar amino acids (**Fig. 2**). Interestingly, several clusters of variants that increased growth rate relative to wild-type due to an apparent positive effect on ADSL function were found, most prominently near the 5ʹ and 3ʹ ends of the gene (discussed later). In a control experiment where ADSL variants were expressed from a weak promoter in the absence of doxycycline (*i.e.* in presence of ADE13), there was no difference between the fitness scores of stop-gain and synonymous variants **(Supp Fig. 2**), indicating that our assay does not discriminate functional from non-functional variants under these conditions. In contrast, when ADSL variants were expressed under a strong promoter in the absence of doxycycline, a slight difference between the effects of stop-gain and synonymous variants could be observed **(Supp Fig. 2**), indicating that high ADSL expression boosts growth alongside ADE13.

### Deriving enzymatic activities from fitness scores

When comparing fitness scores from XS and XL experiments, we observed a nonlinear relationship characterized by a steeper slope at the lower end of the fitness scale (**Fig. 3a**). We hypothesized that this non-linear relationship stems from the inherent non-linearity between growth and enzyme activity as described in the flux control model developed by Kacser and Burns [20]. This pattern arises because the cells are more sensitive to changes in enzymatic activity at low expression levels, where small differences in activity result in relatively larger changes in fitness. Conversely, at high expression levels, the system begins to saturate, leading to diminishing returns in fitness gains despite additional increases in activity. This saturation particularly affects hypomorphic variants, *i.e.,* genes with partial loss of function, as their reduced activity can be compensated by increased expression levels.

**Figure 3:**
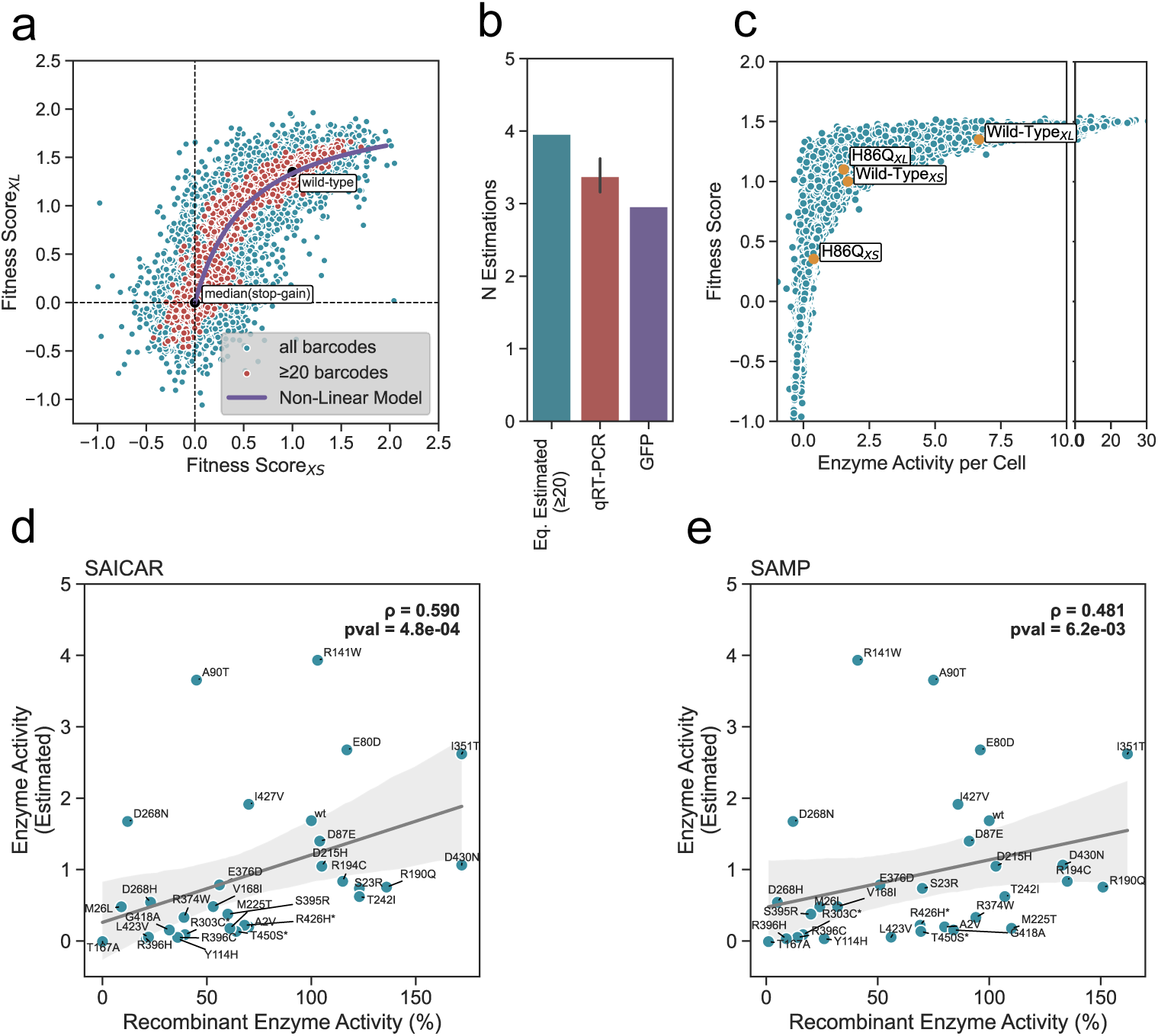
Deriving enzymatic activities from fitness scores via nonlinear modelling. a. Scatter plot of fitness scores from XL (high expression) vs. XS (low expression). Non-linear model fit shown (details in Results). Variants with >20 barcodes are red; others blue. Median scores for stop-gain and wild-type indicated.
b. Validation of model-estimated N (ratio of XL to XS expression) using qRT-PCR and fluorescence (GFP) data.
c. Non-linear relationship between estimated enzyme activity and fitness score. Scores for wild-type and H86Q variants in XS and XL experiments are highlighted.
d. Correlation between model-estimated and recombinant enzyme activities (% of wild-type) for SAICAR catalysis (ρ = 0.59).
e. Correlation between model-estimated and recombinant enzyme activities (% of wild-type) for SAMP catalysis (ρ =0.481). Data from 30 ADSL variants. Variants marked with * have multiple measurements, and the data shown represent the average of these measures.

To study this in more detail, we developed a mathematical model to estimate the enzymatic activity of ADSL variants and relate it to the resulting growth rate (**Fig. 3a**). Our model assumes that each variant *i* is described by a latent parameter, *E_i_*, representing the variant’s specific enzymatic activity (molecules of substrate converted into product per unit time per unit enzyme). Variants can differ in their activity for various reasons, such as differences in thermodynamic stability, which can influence how much properly folded enzyme is available, or differences in the protein’s affinity to its substrate. We also assume that the total enzymatic activity per cell increases by a constant factor, *N*, when expressed from the strong promoter (XL) compared to the weak promoter (XS). For simplicity, we consider that a variant with specific activity *E_i_* has a total enzymatic activity per cell of *E_i_* when expressed from the weak promoter, and N × *E_i_* when expressed from the strong promoter. We then modelled the relationship between enzymatic activity (per cell) and fitness:

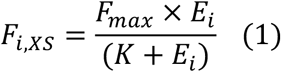

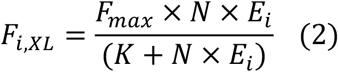

Here, *F_i,XS_* and *F_i,XL_* are the normalized fitness scores for variant *i* measured in the XS and XL experiments, respectively; *F_max_* and *K* are parameters of the model, analogous to the *V_max_* and *K* parameters in the Michaelis-Menten equation; and *N* is the expression level of the XL promoter relative to the XS promoter. After solving (1) for *E_i_* and substituting in (2), we obtained an equation for fitness scores in the XL experiment as a function of scores in the XS experiment:

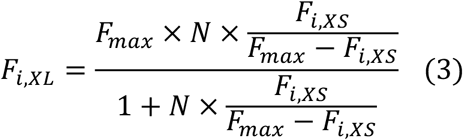

We used equation (3) to derive the values of parameters *F_max_* and *N* by fitting the experimentally measured *F_i,XS_* and *F_i,XL_* scores. The estimated XL promoter strength from the model (N=∼3.96) was similar to values of *N* estimated by flow cytometry (N=∼2.97) and by qRT-PCR (N=∼3.37) (**Fig. 3b**). The model effectively captured the observed nonlinear relationship between fitness scores measured in XS and XL experiments (**Fig. 3a**), and the quality of fit improved when variants were filtered by number of barcodes to reduce experimental noise.

We then derived estimated enzymatic activities for all variants from XS and XL experiments. Variant-specific enzyme activities from the two experiments should be identical; however, due to experimental variance, they exhibit slight differences. We therefore used a weighted mean approach to derive a consensus variant-specific enzyme score based on a combination of XS and XL experiments (see Methods). As expected, low-activity variants experienced a substantial increase in growth when expressed from the strong promoter as compared to the weak promoter (e.g., H86Q, fitness ratio XL:XS=3.11), while highly active variants experienced diminishing returns in growth rate under increased expression (e.g., WT, fitness ratio XL:XS=1.35) (**Fig. 3c**). Estimated enzyme activities became increasingly uncertain for a subset of high-fitness variants, and at the extreme end of the scale; they could not be estimated for variants whose *F_i_* scores exceeded *F_max_* in both the XS and XL experiments. Possible explanations include experimental errors in *F_i_* scores, inaccurate estimation of the *F_max_* parameter, or inadequacy of the model towards the high end of the fitness scale. Nevertheless, estimated enzyme activities were obtained for the vast majority of missense variants (∼92%) with at least one measured fitness score.

We then compared our estimated enzyme activities with those of recombinant ADSL variants, measured as a percentage of wild-type enzymatic activity using biochemical assays. We specifically curated activity data for 30 ADSL variants (A2V, M26L, R141W, R303C, S395R, R426H, T450S, S23R, E80D, D87E, A90T, Y114H, T167A, V168I, R190Q, R194C, D215H M225T, T242I, D268H, D268N, I351T, R374W, E376D, R396C, R396H, G418A, L423V, I427V, D430N, and wild-type) [21]. Our analysis revealed moderate to strong correlations between the estimated and experimentally measured activities, with Spearman correlations of 0.59 for SAICAR and 0.48 for SAMP (**Fig. 3d–e**).

### Mutational constraint at the active site and tetrameric interface

X-ray crystallography shows that ADSL is a homotetrameric enzyme where each monomer comprises 16 α-helices and three small β-sheets, with active sites formed at the interfaces between subunits [22]. Mapping median per-residue enzyme activity scores onto the structure (RSCB-PDB: 2J91), we found that surface residues appeared to be more tolerant of substitutions when compared to buried residues (**Fig. 4a**). When grouping the variants by location, we observed that active site residues were the least tolerant of substitutions, followed by interior and interface residues, while surface residues showed the highest tolerance to substitutions (**Fig. 4b**).

**Figure 4:**
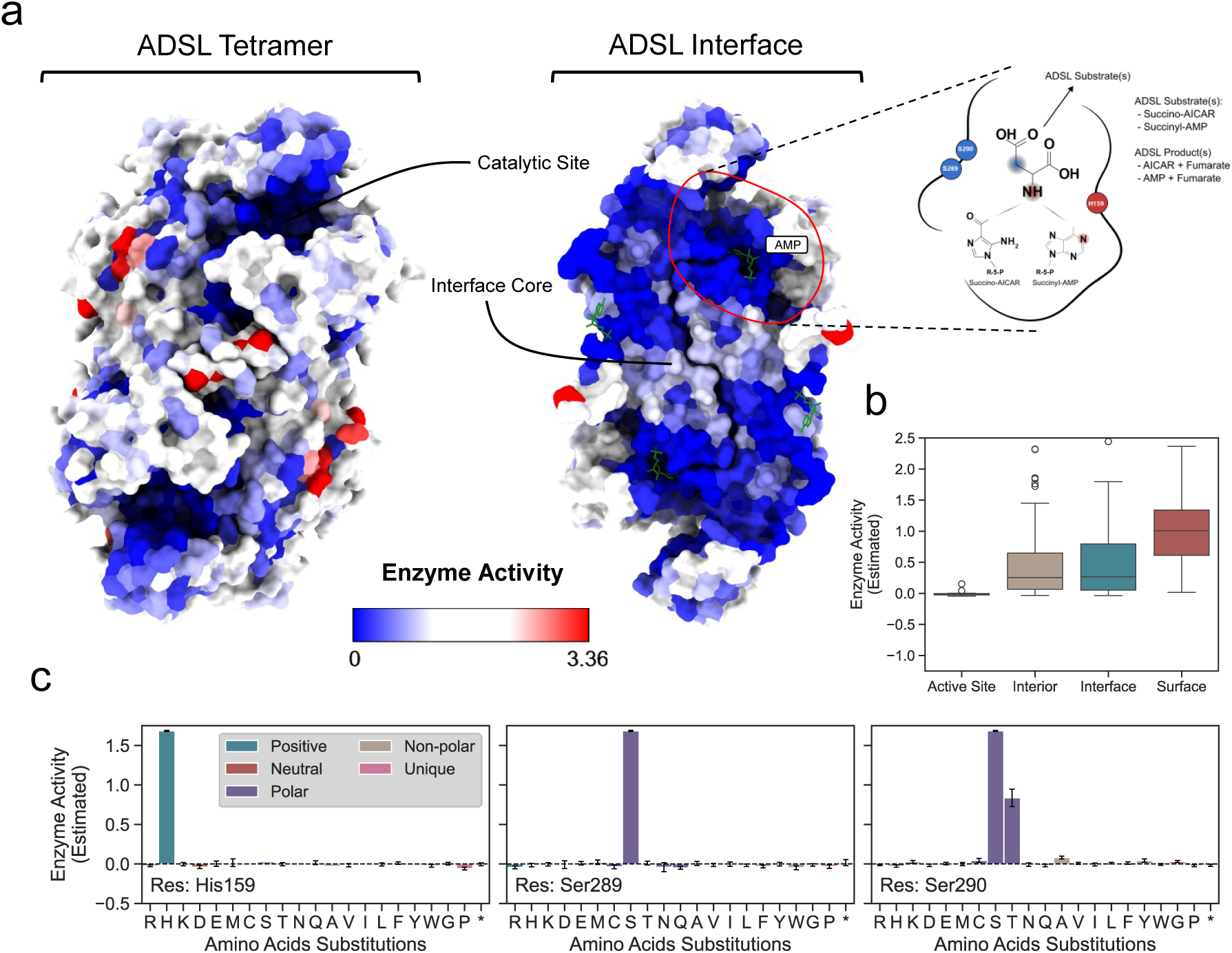
Structural analysis of human ADSL variations via estimated enzyme activities obtained from DMS experiment. a. Estimated enzyme activities mapped onto ADSL crystal structure (PDB ID: 2J91). Left: human ADSL tetramer. Right: ADSL interface with active site and illustration of acid-base catalysis mechanism.
b. Estimated enzyme activities for residues in interior, surface, interface, and active site. Interior and interface residues are generally intolerant, surface residues are tolerant, active site residues are highly intolerant.
c. Estimated enzyme activities for substitutions at catalytic residues His159, Ser289, and Ser290. Substitutions at His159 and Ser289 lose activity; only Ser290Thr retains activity. Error bars show standard error from the model.

We next turned to the catalytic mechanism of ADSL. In the purine biosynthesis pathway, ADSL catalyzes the conversion of SAICAR to AICAR and of SAMP to AMP via a general acid-base catalytic mechanism. Three residues have been directly implicated in catalysis, with His159 protonating the substrate at either the N1 or N6 position to facilitate bond cleavage, and Ser289 and Ser290 thought to initiate the reaction by abstracting a proton from the fumarate moiety of the substrate, though the exact roles of Ser289 and Ser290 are still under consideration due to their positioning [23,24] (**Fig. 4a**). To study this, we analysed the effects of individual missense variants at these three positions by using the estimated enzyme activities obtained from our model. We find that His159, Ser290, and Ser289 are critical for enzymatic activity and are intolerant to most substitutions. Notably, Ser290Thr retains approximately 50% of enzyme activity, while other substitutions reduce estimated enzyme activities to near-zero levels (**Fig. 4c**). This partial retention is likely due to threonine’s structural similarity to serine; both possess hydroxyl (-OH) side chains capable of forming hydrogen bonds and acting as nucleophiles [25].

### Hyper-complementing variants at the N- and C-termini

Variant effect maps showed several clusters of ADSL mutations with hyper-complementing effects on yeast fitness, most notably at the 5ʹ and 3ʹ ends of the gene (**Fig. 5a**). These effects were consistent between XS and XL experiments, reproducible across replicate barcodes, and clustered at specific residues, suggesting a biological basis rather than experimental artefact. Although hyper-complementing variants have previously been associated with deleterious effects in humans [26], their relevance in the context of ADSL remains unclear. Structural mapping did not reveal an obvious mechanism, prompting us to consider effects at the mRNA level.

**Figure 5:**
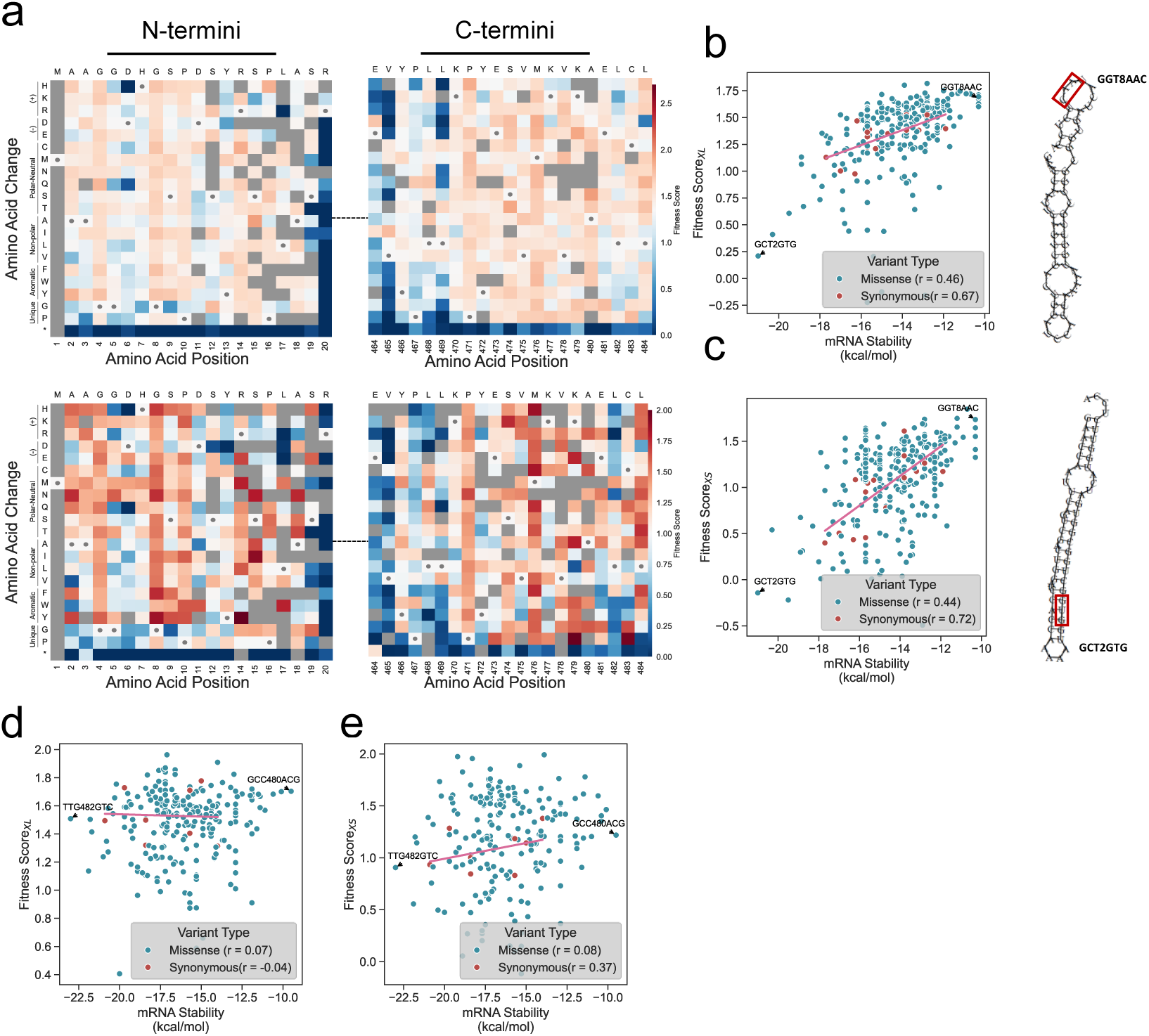
Fitness scores in N- and C-terminus of human ADSL protein. a. Heatmap showing fitness scores in the N- and C-terminal regions from high (XL, top) and low (XS, bottom) expression experiments. Wild-type residues indicated by squares with dots.
b. Correlation between mRNA stability in a 60-nt window around the start codon and fitness scores from XL (high expression) experiment, calculated for N-terminal variants (*codons 1-10*). Least and most stable variants highlighted at the bottom and top, respectively, with 2D mRNA structures showing codon changes (red squares).
c. Same as b, but for XS (low expression) experiment.
d. Correlation between mRNA stability in a 60-nt window around the stop codon and fitness scores from XL (high expression) experiment, in C-terminal variants (*codons 476-485*).
e. Same as d, but for XS (low expression) experiment.

We previously showed that increased stability of mRNA folding around the start codon can lead to dramatic loss of gene expression in bacteria and yeast [27,28]. In agreement with these observations, the predicted stability of mRNA folding in a 60-nt window around the ADSL start codon correlated significantly with the fitness scores of ADSL variants in XS and XL experiments (**Fig. 5b-c**). The wild-type ADSL construct showed relatively strong folding in this region (76^th^ centile of all variants tested when ranked by -ΔG), explaining why most mutations in this region reduced folding and increased growth rate. No comparable effect has been suggested at the 3ʹ ends of human ADSL gene, and mRNA folding at the 3ʹ end showed no correlation with fitness in our data. Therefore, the mechanisms by which C-terminal variants enhance fitness remain uncertain (**Fig. 5d-e**).

### Assessment of variant pathogenicity using DMS measurements

DMS has emerged as a powerful approach for assessing variant pathogenicity, either to aid clinical classification directly [29], or for benchmarking VEP performance [30,31]. While DMS phenotypes may not always fully capture the underlying mechanisms of pathogenicity, top-performing VEPs, such as CPT-1 [32] and AlphaMissense [33], often correlate well with DMS data [34,35]. Notably, our ADSL DMS measurements also show good agreement with VEPs. For CPT, we observe Spearman correlations of 0.660 and 0.663 with XS and XL fitness scores, respectively, which is higher than 30/36 human DMS datasets previously evaluated [36]. For AlphaMissense, we observe similar correlations of 0.663 and 0.670, higher than 33/36 DMS datasets. Interestingly, our estimated enzyme activities show even stronger correlations of 0.694 and 0.699 with CPT and AlphaMissense, respectively, providing further support for the utility of this modelling approach.

To evaluate the performance of our data in classifying clinical ADSL variants, we started by comparing the distribution of estimated enzyme activities across classes of missense variants (**Fig. 6a**). There is a significant difference between pathogenic variants, and those defined as benign, either by having a benign or likely benign classification in ClinVar, or by being present in a homozygous state in gnomAD (p = 0.015, Mann-Whitney U test). However, the variation of scores across both groups is high, with some pathogenic variants having high activity, and one benign variant having close to no activity. In contrast, putatively benign missense variants from gnomAD present in the human population show an even broader distribution, with no significant difference observed when compared to the benign or pathogenic variants. This suggests that there is a considerable presence of damaging ADSL variants present in the healthy population, likely in a heterozygous state.

**Figure 6:**
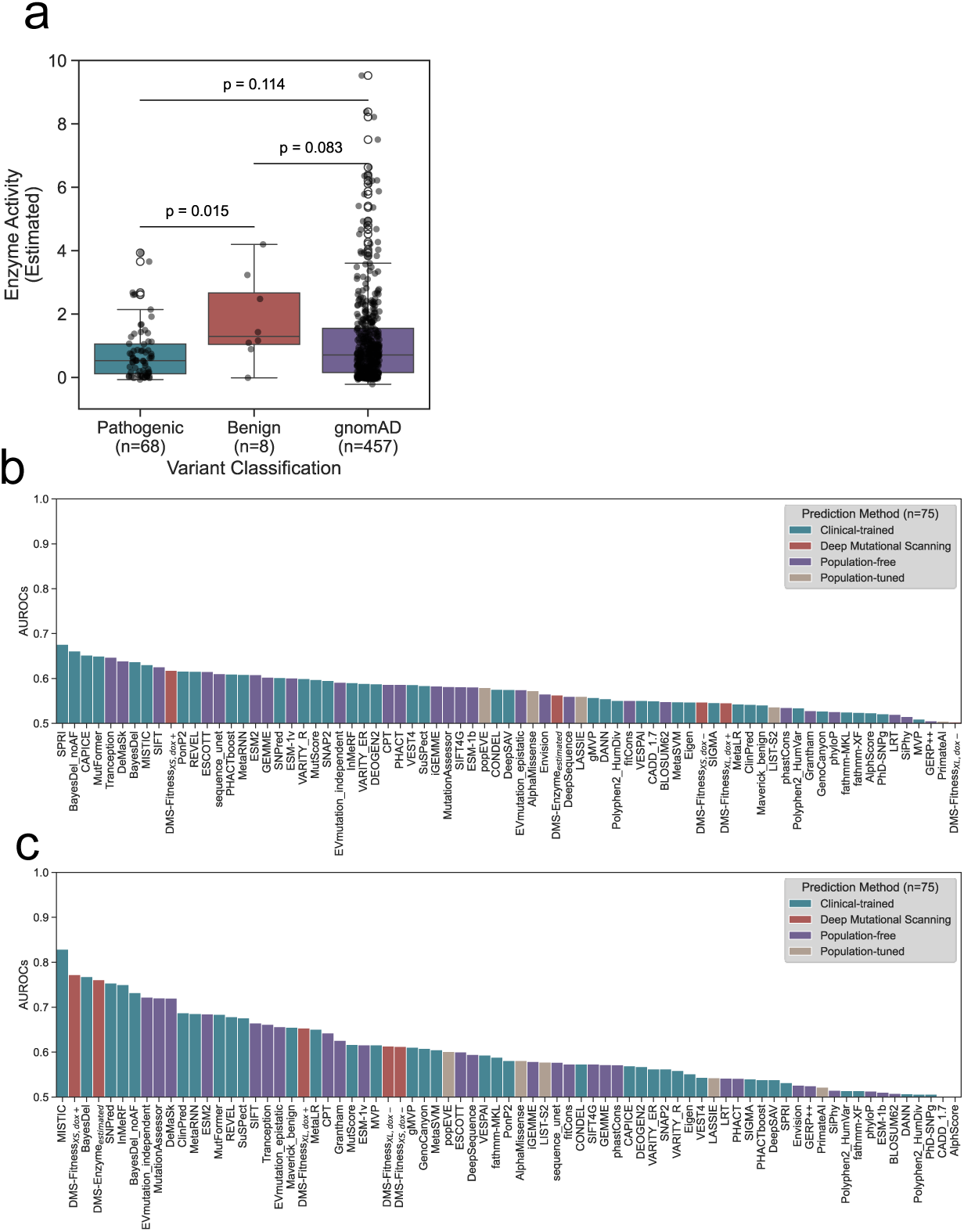
Clinical relevance of DMS-derived scores and VEPs predictions in assessing ADSL variants. a. Box plot of estimated enzyme activities for pathogenic (n=68), benign (n=8), and gnomAD-reported (n=457) missense variants. Pathogenic vs. benign: p = 0.015 (Mann-Whitney U test). gnomAD variants show no significant difference from others.
b. Bar graph showing AUROC values for pathogenic vs. putatively benign variants using DMS-derived scores and VEPs. VEP tools are categorized by training datasets: clinical-trained, population-free, and population-tuned.
c. Same as b, but using pathogenic vs. confirmed benign variants.

Next, we assessed the ability of DMS data to identify pathogenic missense variants, in comparison to 75 different VEPs. The VEPs were classified into three categories based on their potential for inflated performance estimates due to data circularity, as previously defined [36,37]. Circularity arises when predictors are evaluated using data that are not fully independent from their training data, leading to overestimated performance. This can occur at the variant level (e.g. when the same or closely related variants appear in both training and evaluation) or at the gene level (e.g. when variants from the same gene are present in both sets) [38]. *Clinical-trained* VEPs are trained directly on human variants with clinical labels, such as those in ClinVar, and are therefore most susceptible to circularity, especially when assessed using clinical or population data. *Population-tuned* VEPs are not trained on clinically labelled variants but have been exposed to human population data through processes such as calibration, tuning, or the inclusion of allele frequency as a feature. While at lower risk than clinical-trained methods, they may still exhibit bias due to this indirect exposure. Finally, *population-free* VEPs are developed without using any human clinical or population variant data, instead relying on information from sequence conservation, protein structure, or language models. These are not expected to suffer from data circularity when evaluated on human datasets. Although most widely used VEPs to date have been clinical-trained, several recent population-free and population-tuned predictors have demonstrated strong performance while greatly reducing or eliminating the risk of circularity and bias [36,37].

First, we assessed discrimination between pathogenic and putatively benign missense variants, using essentially the same approach that previously highlighted ADSL as a protein-coding gene where VEPs perform poorly [10] (**Fig. 6b**). As expected, all VEPs showed quite low classification performance, with even the top clinical-trained method only achieving an AUROC value of 0.68. However, our DMS data did not improve upon this, with AUROC values of 0.62, 0.55, and 0.56 for fitness scores from XS, XL experiments and estimated enzyme activities, respectively.

The poor performance of DMS data and VEPs in this task presumably reflects the presence of many damaging, potentially pathogenic variants present in a heterozygous state in the human population, suggesting that gnomAD is not an appropriate source of benign variants in this case. Therefore, we also assessed performance at distinguishing between pathogenic and benign missense variants, although the analysis is limited by the fact that the benign set contains only 8 variants. Here, the predictive performance of both DMS data and VEPs improves markedly (**Fig. 6c**). The top-ranking method was MISTIC, a clinical-trained predictor, with an AUROC of 0.83. However, the recent observation that MISTIC apparently performs well in clinical variant classification yet very poorly in terms of agreement with DMS data [36] strongly suggests that its high ranking is likely influenced by circularity due to overlap with training data. Notably, MISTIC heavily relies on allele frequency [39], which is particularly problematic in this comparison, since benign variants are often classified directly using allele frequency as evidence [40]. We also note that MISTIC recently emerged as one of the VEPs showing the highest levels of apparent ancestry bias [37], further suggesting that its apparent performance may reflect overfitting rather than generalisability. Importantly, similar ancestry bias was also observed for the next highest ranking VEPs, BayesDel and SNPred.

Interestingly, DMS data, specifically the XS fitness measurements and the enzyme activity scores, outperform all VEPs other than MISTIC and BayesDel in distinguishing pathogenic from benign missense variants, with AUROC values of 0.77 and 0.76, respectively. While these values still represent far from perfect classification performance, they demonstrate that DMS-based scores can be among the most robust and informative tools currently available for missense variant interpretation.

### Predicting clinical ADSL deficiency using biallelic pathogenicity scores

One limitation of the above approach is that it only considers variants in isolation. However, recessive disease is not caused by individual variants, but instead depends on the combined effects of both alleles together. Given the modest classification performance observed for both VEPs and DMS in single-variant analyses, we wondered whether consideration of the biallelic context, *i.e.,* the variants in both copies of a gene, could lead to improvements.

First, we considered previously published measurements of ADSL enzyme activities in patient fibroblast samples. For each sample, we calculated a biallelic pathogenicity score (BiPS) based on estimated enzyme activities that and compared these values to the conversion rates of SAICAR to AICAR and SAMP to AMP (percentage of healthy individual activity), which were previously measured in patient samples. This analysis revealed a moderate correlation between estimated ADSL activities and those measured in patient samples using the SAICAR and SAMP assays, with Spearman’s correlation coefficients of 0.56 and 0.52, respectively (**Fig. 7a–b**).

**Figure 7:**
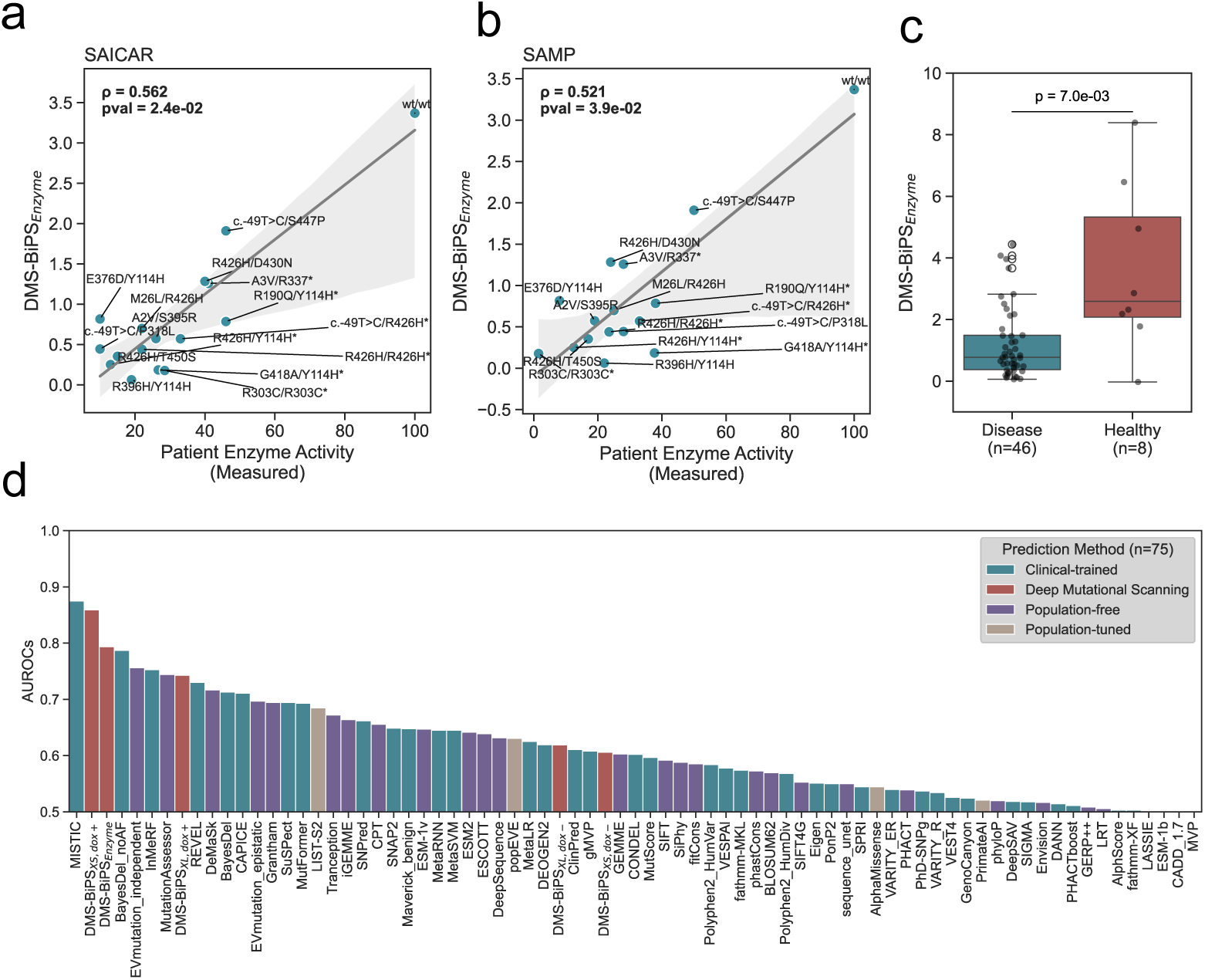
Clinical relevance of biallelic pathogenicity scores in assessing ADSL-D cases and healthy cases. a. Correlation between enzyme activity measured in patient’s fibroblasts and the biallelic pathogenicity score (DMS-BiPS_Enzyme_) for SAICAR catalysis (ρ = 0.56). Dataset is from 16 ADSL patients, with genotypes labeled. Genotypes marked with * include multiple measurements, and the displayed results are the averages of those measurements.
b. Correlation between enzyme activity measured in patient’s fibroblasts and the biallelic pathogenicity score (DMS-BiPS_Enzyme_) for SAMP catalysis (ρ = 0.52).
c. Box plot of biallelic pathogenicity scores (DMS-BiPS_Enzyme_) for disease group (n=46) and healthy group (n=8). Significant difference between groups (p = 0.007, Mann-Whitney U test).
d. Bar graph showing AUROC values for classification performance of biallelic pathogenicity scores from DMS-derived scores and VEPs. Comparison uses 49 genotypes in disease group and 8 cases in healthy group. VEP tools are categorized by training datasets: clinical-trained, population-free, and population-tuned.

Next, we compiled a reference set of 49 distinct genotypes in the disease group, while 8 homozygous healthy genotypes with likely benign or benign variants were compiled as the healthy group. Of the patients, 9 were homozygous for a missense variant, 37 were compound heterozygous for two missense variants, and 3 were compound heterozygous for a missense and a protein null variant. For each disease and healthy genotype, we calculated BiPS by summing the individual scores of each allele. We do this for estimated enzyme activities, fitness scores, and VEP scores. Next, we compared the distribution of BiPS-derived from estimated enzyme activities across variants observed in patients with disease group and healthy group. We observe a clear, highly significant separation between two groups (p = 0.007, Mann-Whitney U test) (**Fig. 7c**). Notably, this separation appears to be markedly stronger than the variant-level analysis in **Fig. 6a**.

Finally, we assessed the predictive performance of BiPS in discriminating between disease and healthy groups, as quantified using AUROC (**Fig. 7d**). Notably, the classification performance is improved across VEPs and DMS compared to the single-variant analyses. As with the pathogenic *vs.* benign variant classification task, the highest AUROC value is observed for MISTIC (0.88), a clinical-trained predictor whose performance is likely inflated by circularity, as previously discussed. However, the XS fitness scores and the enzyme activity scores exceed all other VEPs, with AUROC values of 0.86 and 0.79, respectively. These findings show that incorporating biallelic context enhances predictive power, and that DMS-based biallelic scores offer a functionally grounded and dependable approach for identifying individuals with ADSL deficiency.

## Discussion

Fitness is a common readout in DMS experiments because it is biologically relevant, applicable to many genes, and measurable at scale [26,41,41–43]. However, fitness is often difficult to interpret, as its molecular basis may be unclear and its relationship to underlying biochemical activity is typically nonlinear. To address this, we used a pair of yeast complementation assays performed at different expression levels, allowing us to transform fitness scores into quantitative estimates of enzymatic activity. Compared to fitness, activity scores offer two key advantages: they have a clear biochemical interpretation, and they are additive, making them well-suited for modelling biallelic effects such as compound heterozygosity.

We applied this dual-expression approach to human ADSL, an enzyme involved in purine biosynthesis, using a yeast complementation system. Fitness scores of ADSL variants depended strongly on expression: variants that were severely damaging under low expression became more tolerable when highly expressed. By modelling this relationship, we generated estimated enzymatic activity scores for thousands of missense and stop-gain variants, providing a mechanistically grounded framework for interpreting the functional consequences of ADSL mutations. These results demonstrate how expression-level dynamics can be leveraged to enhance the interpretability of DMS data for metabolic enzymes and potentially other disease-associated genes.

Our analysis assumes that enzymatic activity influences fitness in a nonlinear way. The nonlinear relationship between molecular phenotypes and fitness is well-established in the literature and supported by experimental findings and theoretical models. In their classic paper, Kacser and Burns proposed a framework to explain how changes in activities of individual enzymes impact the overall fitness of an organism, showed how nonlinearity emerges from the interconnectedness of metabolic networks, and provided multiple examples of sigmoidal relationships between enzyme activity and growth [20,44]. Dykhuizen and Dean expanded on Kacser and Burns’s framework by studying *E. coli* under conditions where lactose metabolism limited growth. They showed that permease and β-galactosidase activities directly affect lactose metabolism, but the relationship between these metabolic activities and fitness is non-linear. Their findings revealed a rectangular hyperbolic relationship, where fitness gains taper as enzyme activity nears saturation, aligning with Kacser and Burns’s model [45]. Walkiewicz et al. investigated the in vitro kinetic properties of TetX2 mutants and developed a mathematical model that, based on these physicochemical properties, predicts changes in fitness from biochemical first principles. They emphasized that the model is reversible, enabling the assessment of specific changes in kinetic performance, such as *K*_m_, enzyme expression, and *V*_max_, by analyzing bacterial growth rates [46]. These studies demonstrated that fitness can be leveraged to estimate enzyme kinetics, as enzyme activity directly influences fitness through metabolic pathways, with nonlinear, often sigmoidal, relationships emerging between enzyme performance and growth.

Although we used a Michaelis-Menten-type equation to model the activity-fitness relationship, it is possible that another function would fit the data better. A large-scale yeast study examined how varying the expression of ∼100 genes at ∼100 different levels impacted growth. While some genes exhibited unexpected patterns, enzymes involved in glucose metabolism showed increased fitness with higher expression, consistent with our assumptions [47]. To directly validate the model would require measuring activity of many ADSL variants experimentally, which is currently not possible in a high-throughput format. However, our formulation is supported by the high correlation of predicted enzymatic activities with published data, and by the good fit to the relationship of fitness scores measured in low- and high-expression experiments. Estimated enzymatic activity shows slightly higher correlation with VEPs compared to raw fitness scores, but some enzymatic activity scores are missing, indicating a trade-off between data quality and coverage. We also note that activity does not consistently outperform raw fitness scores in all applications. In particular, the XS fitness data yielded slightly higher AUROC values than the activity scores in the pathogenic vs. benign variant classification task. This may reflect the fact that the activity transformation smooths over some of the more extreme fitness effects, particularly at the low-activity end of the curve, potentially reducing resolution in a classification setting. Nevertheless, the activity scores provide a quantitative, interpretable measure of function that is well suited for modelling variant effects at both the allele and individual level.

Previous efforts to incorporate biallelic context into variant interpretation have demonstrated the value of combining allelic effects for diagnosing recessive disease. In metabolic disorders such as phenylketonuria, the Genotypic Phenotype Value (GPV) model has successfully stratified clinical severity by scoring each allele independently and integrating the scores to predict patient phenotype [48]. Other computational frameworks, including HMMvar and MOI-Pred, have highlighted the importance of modelling compound heterozygosity by quantifying joint variant effects [49] or predicting mode of inheritance directly from variant-level features [50].

More recently, experimental approaches have also moved toward biallelic modelling: in recessive muscular dystroglycanopathies, DMS data have been used to derive additive scores from dual alleles, correlating with age of onset and disease severity [51]. Gene-level pathogenicity scores such as GenePy have further demonstrated how aggregate mutational burden can help prioritize recessive diagnoses in large cohorts [52]. Our BiPS model builds on these efforts by directly linking mechanistically interpreted enzymatic activity scores to clinical genotype–phenotype relationships in ADSL deficiency. Unlike previous models that rely on empirical genotype data or computational heuristics, BiPS integrates experimental measurements of enzyme activity from both alleles to generate a quantitative prediction of disease likelihood. This dual-layered framework—combining functional modelling with allelic integration—improves diagnostic classification and provides a generalizable strategy for variant interpretation in autosomal recessive disorders.

Our analysis reveals a group of hyper-complementing variants that exhibit higher functional scores than the wild-type gene. Some of these effects can be explained by reduced mRNA folding energy around the ADSL start codon, which is expected to enhance translation efficiency. This translation-enhancing mechanism has been observed in bacteria and yeast but not in humans [27,28,53], making it a potential source of bias in yeast-based MAVEs. In contrast, hyper-complementing variants near the 3ʹ end of ADSL could not be explained by mRNA structure, and the molecular basis of these effects remains unclear. While it has been proposed that hyperactive mutants may go undetected due to their subtle phenotypic effects [44], deep mutational scanning frequently reveals such variants, albeit typically in limited regions or specific residues. For example, PAX6 Ile71 variants were recently shown to enhance DNA binding to a specific regulatory element [54], and INSR G333 variants increased insulin binding affinity [55]. Hyperactive variants are often deleterious in humans [26], and the ability of MAVEs to detect these gain-of-function effects is a key advantage over current variant effect predictors (VEPs), which typically lack sensitivity to increased activity. Notably, recent studies have implicated increased ADSL activity in cancer via modulation of the STING and autophagy pathways [56,57], raising the possibility that some of the gain-of-function variants identified here may play a role in tumorigenesis.

## Materials and Methods

### Strain preparation and yeast complementation

We purchased the yTHC-ADE13 strain from the Yeast Tet-Promoters Hughes Collection, pADE13: kanR-tet07-TATA URA3: CMV-tTA MATa his3-1 leu2-0 met15-0. In that strain, the expression of ADE13, the yeast homologue of human ADSL, is controlled by the tet07 promoter. When exposed to doxycycline treatment (10-20 µg/mL), the yTHC-ADE13 strain suppresses ADE13 expression. To express the human ADSL gene in this strain, we employed Gateway-compatible destination vectors named pMP1_XS and pMP1_XL. These vectors include a CEN/ARS-based element, yeast expression promoter, and a HIS3 marker. The human ADSL cDNA construct (UniProtKB: P30566) was designed using the IDT codon optimization tool to align with the codon preferences of *Saccharomyces cerevisiae (S. cerevisiae)*. The ADSL cDNA was then cloned into the Gateway Entry vector pAINt and then introduced into the pMP1_XS and pMP1_XL plasmids through a Gateway LR reaction. Subsequently, the resulting constructs were transformed into the parental yTHC-ADE13 strain using LiAc/SS carrier/PEG method [58]. Transformants for XS and XL were grown for 48 hours, with the media refreshed every 12 hours, during which doxycycline was administered to suppress ADE13 expression. After the suppression of ADE13 expression, the optical density at 600 nm was measured every 15 mins for approximately 24 hours using Tecan Infinite M200 Pro.

### Logistic growth model and growth rate calculation

We calculated growth rates via a logistic (sigmoidal) growth model [59]. The standard form of logistic equation (Eq. 1) is fitted to optical density measurements to calculate growth rates and growth rates are used to normalize fitness scores measured in XL and XS experiments.

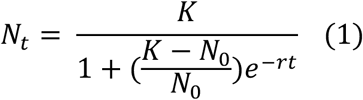

where N_t_ represents the population size at time *t*, number of cells at time *t*, N_0_ denotes the initial population size, K is the carrying capacity, and r is the intrinsic growth rate, which reflects the rate of growth in the absence of any limitations on the total population size.

### Library preparation and barcode variant matching

To generate the ADSL library, we used the plasmid-based saturation mutagenesis protocol originally developed by Wrenbeck et al. [60], using pAINt-ADSL as the template for mutagenesis. This protocol enables the generation of a library that encompasses every possible missense and stop gain variants using mutagenic oligos containing a central degenerate NNS codon. We created the library in 24 segments, each consisting of 20 amino acid positions. The barcode-based approach is helpful in deep-scanning experiments as it mitigates the impact of sequencing errors, reduces costs, and provides internal replicates, enabling the differentiation between low- and high-quality variants [43]. In our study, each variant in the library was assigned a unique 30-nt barcode. To match barcodes with ADSL variants, called library phasing, we used the PacBio Sequel IIe sequencing system. The system’s HiFi CCS (Circular Consensus Sequencing) reads produce highly accurate sequencing data for long sequences, which includes a unique barcode, a promoter region, and ADSL cDNA sequence containing a variant in our case.

We employ the alignparse Python package to align the reads to the target amplicon and call consensus sequences based on the relevant CCS reads [61]. Following alignment and parsing, a filtering process is applied to remove low-quality reads based on error rates (1-accuracy) with a cut-off of 1e-4. A consensus sequence is then generated for each barcode: if there is only one CCS per barcode, that sequence becomes the consensus; if multiple CCSs are present, they are combined to build the consensus. However, if differences or high-frequency non-consensus mutations are detected between CCSs associated with the same barcode, the entire barcode is discarded to ensure accurate barcode variant matching.

### Competitive growth assay

We introduced pMP1_XS-ADSL and pMP1_XL-ADSL libraries into the yTHC-ADE13 strain using the LiAc/SS carrier/PEG method for high efficiency yeast transformation [58]. In each library, a minimum of one million yeast cells were transformed. Following transformation, yeast cells were allowed to recover for 24 hours in 500 mL of Synthetic Complete Medium without Histidine (SC-His, Formedium) without doxycycline (Sigma-Aldrich) treatment. Following the recovery period, we applied a treatment of 20 µg/mL doxycycline to suppress the yeast ADE13 gene expression, thereby revealing the phenotypes of human ADSL variants within cells. We cultivated the culture for 10 time-points by refreshing the media and sampling every 12 hours. Control experiments were performed without doxycycline treatment.

Subsequently, we did yeast-plasmid extraction by resuspending yeast pellets in P1 buffer from the Qiagen MiniPrep Kit, treating the cells with DTT (Qiagen) and Zymolyase 20T (MP Biomedicals), and then continuing with the standard Qiagen MiniPrep Kit protocol. Following plasmid extraction, we amplified the barcodes using Phusion® High-Fidelity PCR, running 15 cycles to minimize PCR biases. Forward primers are indexed to label conditions and time-points in competition experiment. The PCR products were then separated and collected using E-Gel™ SizeSelect™ II Agarose Gels, 2% (Thermo Fisher Scientific).

### Sequencing and fitness score calculations

To assess the enrichment scores of each variant, we used the Illumina NextSeq 2000 system to sequence the barcode libraries. We used the Enrich2 pipeline developed by the Fowler group for variant scoring using sequencing data derived from DMS experiments [62]. Enrich2 counted barcodes for each condition and time point, including only those with more than 10 counts at time point 0 to avoid processing low-quality reads. We generated scores per-barcode using Enrich2 and calculated variant scores by taking the median of barcodes assigned to each variant and their standard error of the mean. To make fitness scores estimated in the XS and XL experiments directly comparable with each other, we rescaled the data to fit within a range of 0 to 1 in XS and 0 to 1.35 in XL datasets, to match the relative growth rates (fitness) of pMP1-XS-ADSL and pMP1-XL-ADSL strains quantified by growth curve measurements as described above. Specifically, we set the median fitness scores as follows: fitness score (XS: stop gain) = 0, fitness score (XL: stop gain) = 0, fitness score (XS: wild type) = 1, and fitness score (XL: wild type) = 1.35.

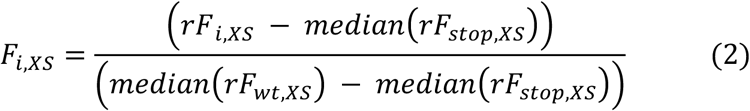

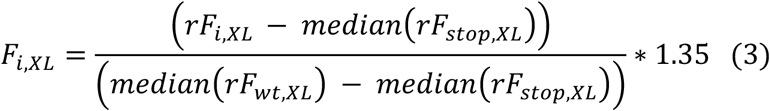

rF_i,XS_ and rF_i,XL_ represent the raw fitness scores for variant i in these experiments. Similarly, rF_stop,XS_ and rF_stop,XL_ are the raw fitness scores for stop gain variants, and rF_wt,XS_ and rF_wt,XL_ are the raw fitness scores for the wild type in the XS and XL experiments.

To optimize the data-point selection used in variant scoring process, we calculated the ratio of the mean variation of fitness scores of barcodes corresponding to the same variant to the variation across the entire set of barcodes, which gives us the experimental error metric (Eq. 4). A lower value of experimental error metric indicates less noise relative to the signal. In this way, the equation aids in identifying the set of time-points with the highest signal-to-noise ratio.

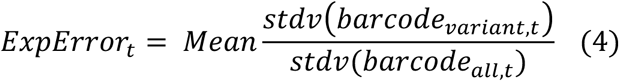

where ExpError_t_ represents the experimental error at certain time point, barcode_variant,t_ refers to the barcodes assigned to the same variant, and barcode_all,t_ denotes all barcodes scored at certain time point. With this approach, we selected time points 0-7 (0-96 hours) for analyzing XS and XL data. Including later time points led to increased experimental error, likely due to challenges in fitting a reliable regression line (refer to the Enrich2 methodology for details).

### Estimation of enzyme activities from fitness scores

We used fitness scores F_i,XS_ and F_i,XL_ to model enzymatic activities of each variant, E_i,XS_ and E_i,XL_, as follows:

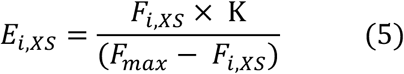

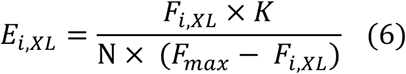

In these equations, *F_i_*_,*XS*_and *F*_i,*XL*_are variant-specific fitness scores measured experimentally, and N = 3.96 and *F_max_* = 1.57 are parameters of the model, derived as described in the results section. K is a scaling factor that cannot be derived from the data and is set to K=1.

We then modelled the standard deviations (SD) of barcode fitness scores using a polynomial fit of SD against fitness scores, and calculated standard errors of fitness score, SE(*F_i_*_,*XS*_) and SE(*F_i_*_,XL_) by dividing the fitted SD by the square root of the number of barcodes. We then used error propagation to derive standard errors of enzymatic activities:

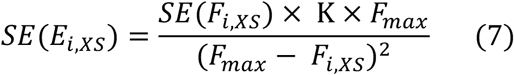

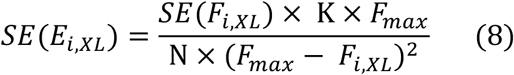

The variant-specific enzymatic activities *E_i,XS_* and *E_i,XL_* are expected to be invariant with respect to expression level. To aggregate the two enzymatic activity scores into a single score, E_i_, we calculated E_i_ as a weighted mean of E_i,XS_ and E_i,XL_ :

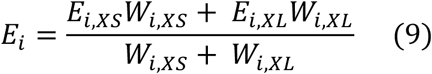

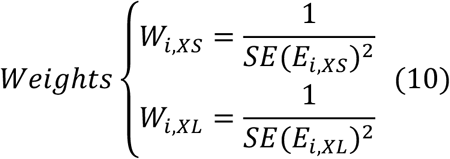

The weights W_i,XS_ and W_i,XL_ are calculated using the inverse-variance weighting method, which assigns higher weights to values with smaller standard errors and lower weights to values with higher standard errors (Eq. 10). Missing data were handled appropriately to ensure that only valid data contributed to the final calculations.

### Structural analysis

We calculated the location of residues in the protein by considering a threshold of 25% relatively accessible surface area (RSA) to assign residues a classification of the interior (below threshold)/surface (above threshold). The fitness scores were used to replace the B-factor values in the PDB file by calculating the median scores at the corresponding positions, by excluding stop-gain variants. This provides a simple method for projecting median fitness scores per residue onto the protein structure. We used the ChimeraX software to visualize map scores to an experimentally determined crystal structure of the ADSL tetramer (RSCB PDB ID: 2J91). Secondary structure is visualized using the SSDraw Python package [63], with evolutionary conservation calculated from the ConSurf Server integrated [64].

### Collection of variant and patient datasets and classification performance

To collect clinically reported benign and pathogenic ADSL variants, we used ClinVar (2024.12.05) and case reports published (2025.05.03). 73 missense variants in the human ADSL gene have been reported with a likely pathogenic or pathogenic phenotype, while 6 missense variants have been reported with a benign or likely benign phenotype, including two homozygous missense variants from gnomAD, resulting in a total of 8 benign variants. We also considered missense variants reported in gnomAD v4.1 as putatively benign, collecting 554 missense variants from the database **(see Supp. Table 1**).

We compiled clinically reported ADSL deficient cases into the “disease group” **(see Supp. Table 2**), including missense and stop-gain variants, while removing duplicates to retain unique entries. This resulted in 49 distinct genotypes in the disease group, while 8 homozygous healthy genotypes with likely benign or benign variants were compiled as the healthy group. We gathered enzyme activity data, as a percentage of normal (wild-type) activity, from patient fibroblast samples **(see Suppl. Table 3)** and from recombinant enzyme tests provided by Marie Zikanova, Vaclava Skopova, and Stanislav Kmoch, as well as from a review of existing studies **(see Suppl. Table 4)**.

VEPs were selected based on those used in a recent benchmarking study [36]. We filtered for coverage in ADSL, considering only those 75 VEPs with scores available for at least 85% of pathogenic variants, 85% of benign variants, and 85% of putatively benign variants. AUROC values were calculated using the roc_auc_score() function from the sklearn.metrics module in Python, as implemented in the custom roc_auc_calculate() function.

### Biallelic pathogenicity scores

An additive model integrates the contributions of both alleles to the overall phenotype, combining their estimated effects into a single Biallelic Pathogenicity Score (BiPS) (Eq. 11). We derived BiPS from fitness scores, estimated enzyme activities, and VEP scores and assessed its effectiveness in classifying clinically reported cases.

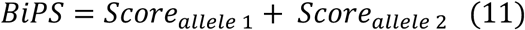

## Data availability

Varant scores will be deposited in MaveDB, sequencing data in GEO, and VEP scores in OSF or figshare prior to publication.

## Acknowledgments

We thank Ben Livesey for providing VEP scores.

## Declaration of generative AI and AI-assisted technologies in the writing process

During the preparation of this work the authors used the AI tool ChatGPT in order to improve language and readibility. After using this tool/service, the authors reviewed and edited the content as needed and take full responsibility for the content of the publication.

## Supplementary Tables

**Supplementary Table 1:** Human ADSL variants from ClinVar database, including putatively benign variants from gnomAD and those identified in case reports.

**Supplementary Table 2:** Unique genotypes of ADSL Deficiency cases and healthy individuals from case reports and database: www.adenylosuccinatelyasedeficiency.com.

**Supplementary Table 3:** Enzyme activities measured in patient fibroblasts from ADSL deficient patients.

**Supplementary Table 4:** Recombinant enzyme activity of human ADSL variants.

---

**Supplementary Figure 1:**
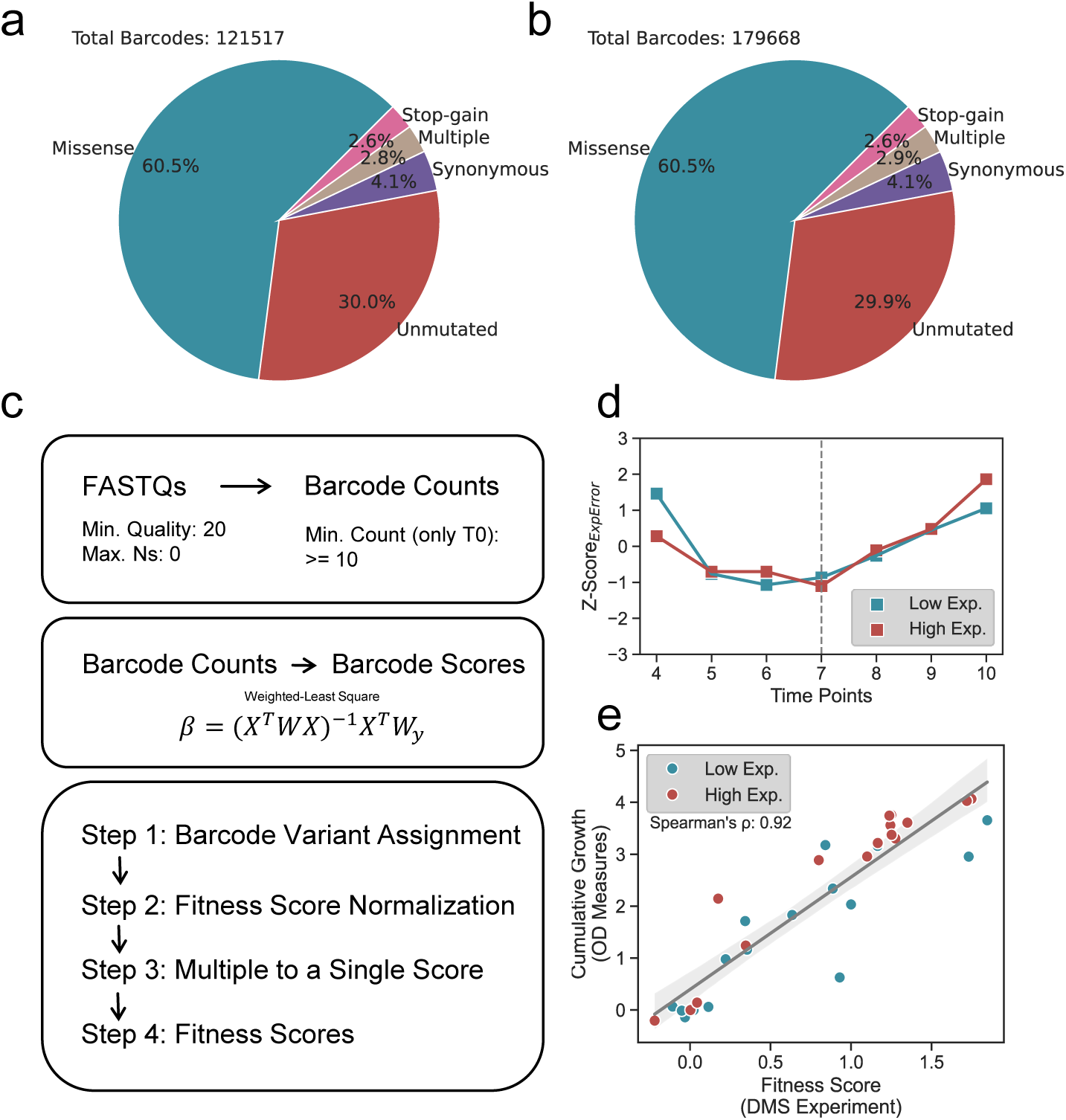
Analysis of human ADSL library and growth-based competitive assays. a. Distribution of barcodes in the XS library: approximately 60% missense, 30% wild-type, remainder stop-gain, multiple, and synonymous variants. Total barcodes: 121,517.
b. Same as a, but for the XL library. Total barcodes: 179,668.
c. Schematic representation of variant scoring process: barcode counting, per-barcode scoring, and score extraction. Quality control: sequencing quality ≥20, barcodes ≥10 counts at time point 0.
d. Plot of Z-scores for experimental error at different time points for XS and XL experiments. Minimal error observed at time point 7 for both conditions.
e. Scatter plot showing correlation between normalized fitness scores from deep mutational scanning (DMS) and cumulative growth from optical density-based growth assays for 14 ADSL variants. Spearman’s correlation coefficient: 0.92.

**Supplementary Figure 2:**
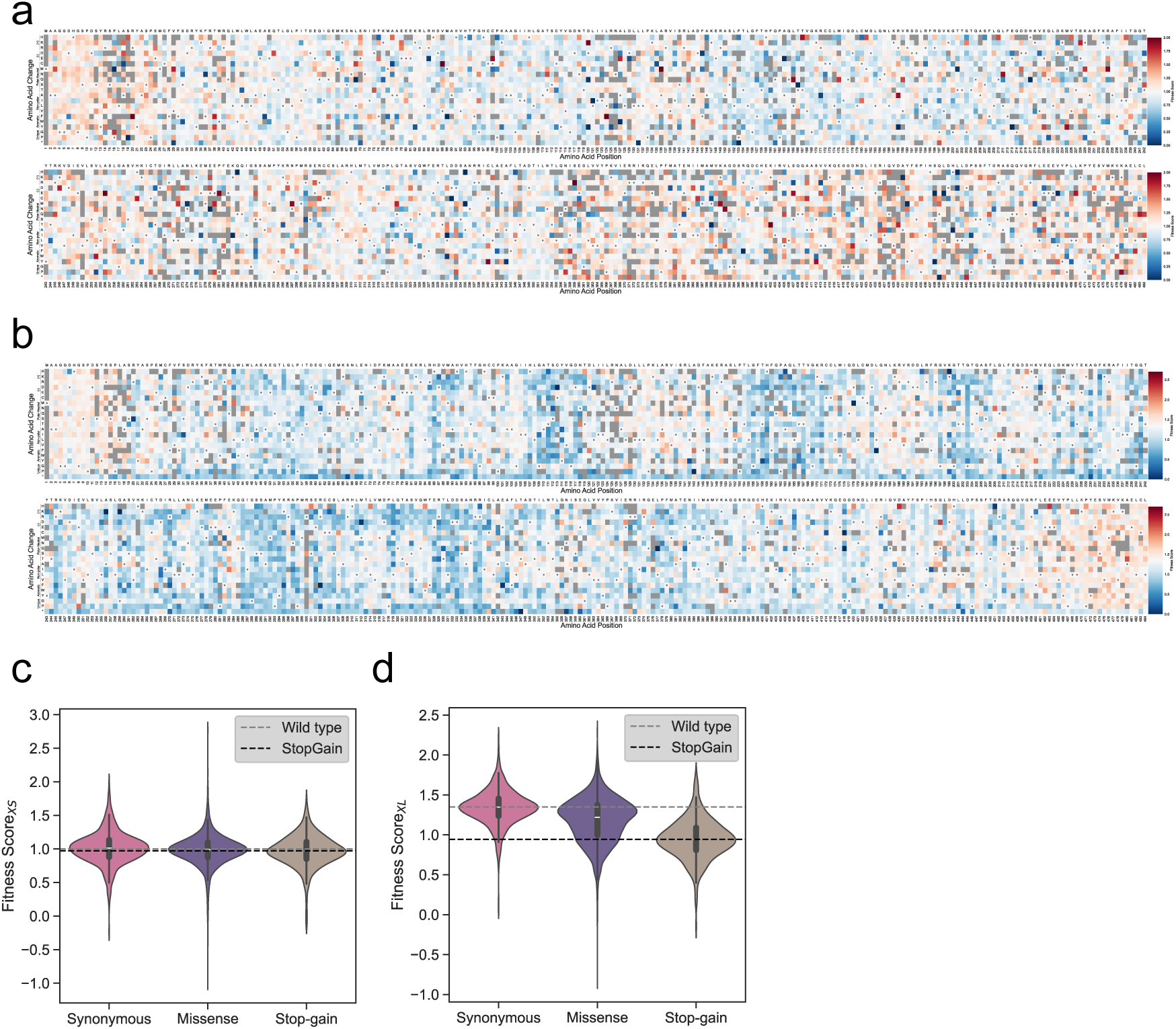
Fitness score landscape for the co-expressed ADE13 and human ADSL gene under low expression and high expression conditions. a. Heatmap of fitness scores from the XS experiment without doxycycline treatment. Color key: darker blue for damaging variants, white for wild-type-like, red for hyper-complementing variants.
b. Same as panel a, but for the XL experiment.
c. Distribution plot of fitness scores for the XS experiment without doxycycline treatment, showing scores for stop-gain, missense, and synonymous variants. Median scores for wild-type and stop-gain variants are marked with dashed lines.
d. Same as panel c, but for the XL experiment. In this panel, stop-gain variants show a distribution shifted towards lower fitness scores compared to synonymous and most missense variants.

